# Performance and impact of using a rapid molecular test to detect *Chlamydia trachomatis* and *Neisseria gonorrhoeae* in women suspected of having pelvic inflammatory disease

**DOI:** 10.1101/2020.10.23.351825

**Authors:** J Munrós, A Vergara, E Bataller, G Restovic, B García-Lorenzo, MJ Álvarez-Martínez, A Mira, F Carmona, J Vila, J Bosch

## Abstract

**Objective:** The diagnosis of pelvic inflammatory disease (PID) is challenging. Testing for *Chlamydia trachomatis* (CT) and *Neisseria gonorrhoeae* (NG) in the lower genital tract is recommended, since a positive result supports the diagnosis. The aim of this study was to investigate the prevalence of CT/NG infection in women suspected of having PID and the usefulness of a rapid molecular test to detect CT/NG.

**Methods:** This observational study included 3 groups of patients: mild-to-moderate PID (n=33), severe PID (n=29) and non-specific lower abdominal pain (NSAP) (n=13). CT/NG infection were analyzed using a standard and a rapid test. A cost analysis was carried out.

**Results:** The presence of CT/NG was determined in 75 endocervical and urine samples. Endocervical samples of 19 patients (25.3%) were CT/ NG positive (two cases of co-infection). NG was not detected in urine in one case. Concordance between rapid and standard tests was 100%. However, the mean time to achieve results was shorter with the rapid test: 2.22 *vs.* 24.37 hours, respectively (*p* < 0.001). No significant differences were observed in the presence of CT/NG in mild-to-moderate compared to severe PID. Costs differed according only to disease severity but to the presence of CT/NG. Only one patient with NSAP was positive for CT.

**Conclusions:** Rapid molecular tests could help with the diagnosis of PID in sexually active women in clinical settings in which a standard technique is not available. Nonetheless, a positive test for CT/NG may not be determinant of the clinical management. The only cost difference relates to disease severity.

## INTRODUCTION

Pelvic inflammatory disease (PID) is a clinical syndrome of the upper female genital tract which is mainly due to polymicrobial infection ascending from the endocervix^1,2^. PID may be also a sexually transmitted disease (STD), with *Neisseria gonorrhoeae* (NG) and *Chlamydia trachomatis* (CT) being the most common causal agents. Other pathogens causing PID are *Mycoplasma genitalium* (MG) and microorganisms from the vaginal microbiota^1,3–6^.

The diagnosis of PID may be challenging because of the lack of definitive diagnostic criteria^7^. Moreover, patients may report a wide range of clinical manifestations, from mild symptoms to very severe disease^1,2,8,9^, which may be confounded with other etiologies, leading to misdiagnosis and treatment delay and short-(tubo-ovarian abscess and pelvic peritonitis) or long-term complications (impaired fertility, ectopic pregnancy and chronic pelvic pain)^1^. Furthermore, PID represents a significant economic burden considering its management and follow-up and possible long-term complications^4,7,10–13^. Therefore, guidelines strongly recommend the initiation of presumptive medical management and early antibiotic treatment in suspected PID cases^2,14,15^.

Testing for CT and NG in the lower genital tract is recommended, and while their absence does not exclude PID, a positive result would support the diagnosis^1,2,5,9,14,15^. However, standard methods for determining the presence of CT and NG are not available in all clinical settings, and are costly and require a lengthy period from sample collection to result obtainment, thereby limiting their use in the clinical management of PID^1,16^. Therefore, the use of a rapid molecular test for CT and NG detection could help in the diagnosis and management of this gynecological pathology. Results would be available more rapidly to take medical decisions for the clinical management of patients with suspected PID compared with the standard methods.

The aim of this study was to investigate the prevalence of CT and NG infection in women suspected of having PID and the usefulness of a rapid molecular test to detect CT and NG for the diagnosis and clinical management of PID.

## MATERIALS AND METHODS

### Study design and subjects

In this observational study, 75 women with suspected PID or non-specific lower abdominal pain (NSAP) were prospectively recruited in a tertiary health care center from April 2016 to April 2017. The study was approved by the Ethics Committee of the hospital (HCB/2016/0171). All women provided written informed consent.

All women meeting clinical criteria for PID diagnosis according to international guidelines^2,14,15^ or who referred NSAP in the emergency department (ED) were asked to participate. The following epidemiological data were recorded: age, tobacco use, previous STD and number of pregnancies.

Patients were managed according to the protocol of the Gynecology Department for the management of PID based on international guidelines^2,14,15^. The clinical criteria for initiating presumptive treatment for PID in a sexually active woman at risk of STD who referred pelvic or lower abdominal pain were cervical motion tenderness or uterine or adnexal tenderness. The following complementary tests were performed in women fulfilling the clinical criteria for PID: blood analysis to determine white blood cell count (WBC), prothrombin time (PT), and C reactive protein (CRP) and screening for other STDs (human immunodeficiency virus, hepatitis B and syphilis); vaginal and urine cultures, endocervical and urine tests for CT and NG, and transvaginal ultrasonography (TUS). Complementary imaging tests were performed if considered clinically necessary. Pregnancy tests were also performed in premenopausal women. The following criteria were used to support the diagnosis: body temperature >38 °C, mucopurulent cervical discharge or cervical friability and elevated CRP value.

Patients were classified as having mild-to-moderate or severe PID according to clinical symptoms and signs and complementary test results^3,17,18^. Severe PID was defined in this study as the presence of severe symptoms or signs and/or the presence of tubo-ovarian abscess at imaging tests^3,17^. Patients who didn’t meet these criteria were classified as mild-to-moderate PID^17,18^. Patients with severe PID were admitted to hospital, while mild-to-moderate cases were managed as outpatients in cases of non-pregnant women with the absence of severe clinical manifestations, nausea/vomiting and comorbidities^1,3,14,17,18^.

The antibiotic regimen was chosen according to the hospital protocol based on local antimicrobial sensitivity (Appendix I). Surgery was considered in cases of diagnostic uncertainty or severe cases presenting parenteral treatment failure.

Follow-up was performed 6 weeks after diagnosis to evaluate clinical symptoms and patient improvement. CT and NG diagnosis tests were repeated in patients previously positive for CT and/or NG.

Additionally, all women referring NSAP in the ED, in whom other causes for these symptoms were excluded but did not meet all PID criteria^2,14,15^, were also invited to participate in the study. In these cases, only blood analysis for WBC, PT and CRP evaluation, and urine test and endocervical swab for detection of CT and NG were performed. Analgesic treatment was provided, if necessary. Follow-up was also performed 6 weeks after diagnosis. If CT and/or NG were detected, the women were promptly informed and appropriate treatment was indicated.

### Sample collection and analysis

Endocervical samples were collected with nylon swabs for the detection of CT and NG. Urine samples were collected for culture in cystine–lactose–electrolyte-deficient agar and for the detection of CT and NG. Gram-stained vaginal smears were analyzed to evaluate potential bacterial vaginosis using the Nugent score.[19] Additionally, intraoperative intraabdominal fluid samples from patients undergoing surgery were cultured.

### Molecular pathogens detection

DNA from endocervical and urine samples was extracted with the Biorobot EZ1® (Qiagen, GmbH, Hilden, Germany). CT and NG were detected with real-time polymerase chain reaction [Anyplex® CT/NG Real-time Detection kit (Seegene, Seoul, Korea)] using a real-time thermal cycler, SmartCycler® (Cepheid, Sunnyvale, California, US). The same samples were used to directly detect CT and NG, without previous DNA extraction, with the GeneXpert® CT/NG assay (Cepheid, Sunnyvale, California, US). Molecular tests for CT and NG were performed immediately after collection with GeneXpert® or three times per week (with refrigeration for less than 48 hours) with Anyplex®.

The conventional and GeneXpert® results were compared, and the time in hours from sample reception at the Microbiology Laboratory until result obtainment was registered for both methods. Samples and DNA were stored at −20°C for further analysis. *Mycoplasma genitalium* and *Trichomonas vaginalis* were retrospectively tested, with the RealCycler® Monotest MGTVUS (Progenie, Valencia, Spain) in a SmartCycler®.

### Cost analysis

Based on the results and information obtained from the clinical study, a cost study was carried out^20^ considering only direct costs. The economic variables included the type and frequency of resources used by each patient including: diagnostic tests, pharmacologic treatments, need for hospitalization, visits, diagnostic imaging tests and laboratory tests. Unit costs in Euros 2018 for each resource used were obtained from the hospital administrative database.

The mean cost per patient was computed using individual patient data. Moreover, cost by disease severity (NSAP, mild-to-moderate PID, severe PID), and the presence of CT and/or NG (at least one positive *vs.* both negative) was also calculated.

### Statistical analysis

Statistical analysis was performed with the Statistical Package for the Social Sciences software, v20.0 for Windows (SPSS, Chicago, Illinois). Continuous variables were compared using the parametric Student’s T-test or one-way ANOVA with the Bonferroni post hoc test and presented as mean and standard deviations, or the nonparametric Mann-Whitney U test and presented as median with interquartile range (ie. 25^th^-75^th^ percentiles). Categorical variables were compared using the Chi-squared test or Fisher’s exact test and presented as total count and relative percentages (%). Statistical significance was defined as a p-value <0.05. Cohen’s kappa coefficient was calculated to assess concordance between the laboratory methods to detect CT/NG.

## RESULTS

### Characteristics of the study population

A total of 75 patients were included in the study and classified into three groups: mild-to-moderate PID (n=33), severe PID (n=29) and NSAP with no clinical suspicion of PID (n=13).

Table 1 shows baseline characteristics, findings at physical examination, laboratory tests and TUS of the patients included. All pregnancy tests performed in women of reproductive age were negative as was screening for other STDs in all the patients included.

**Table 1.**
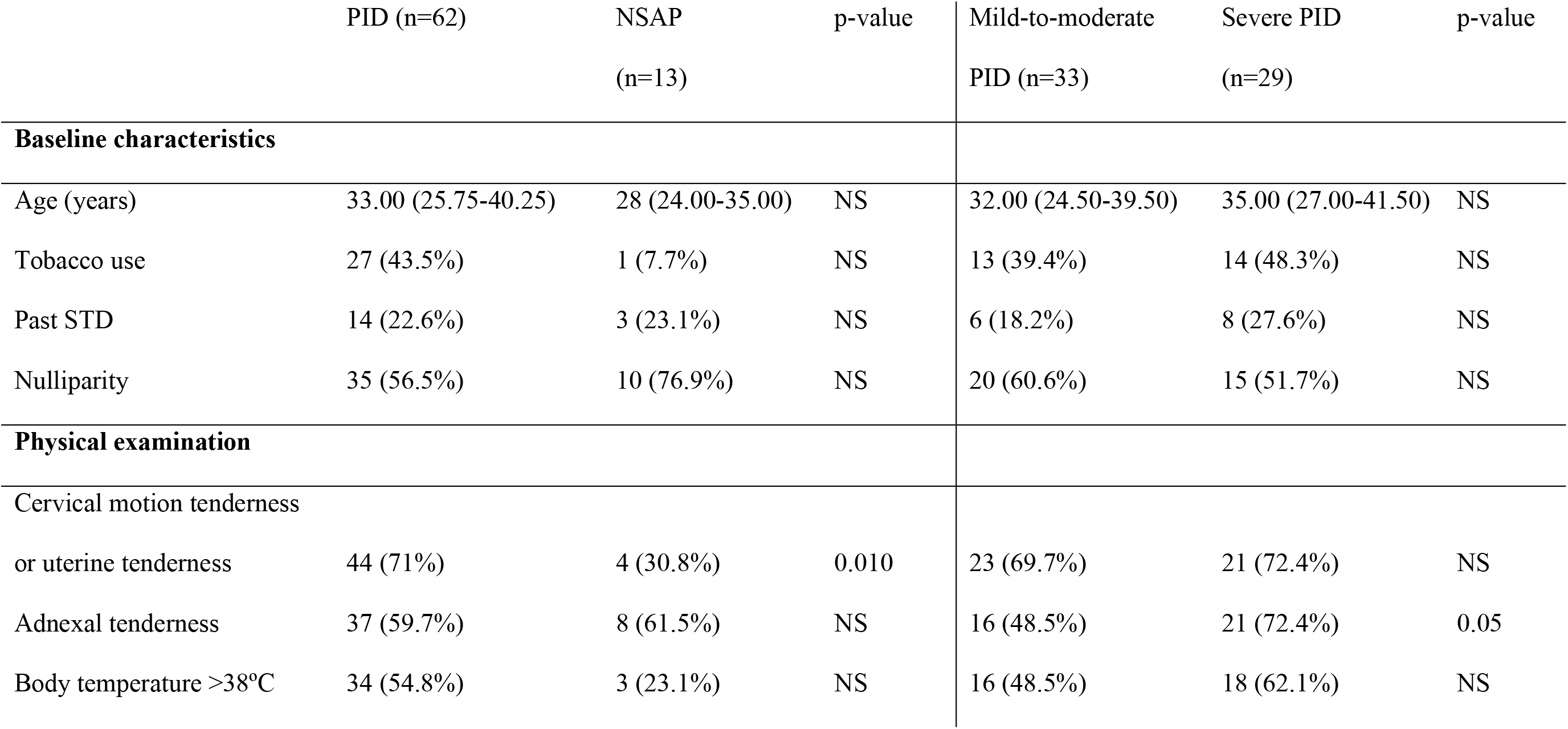
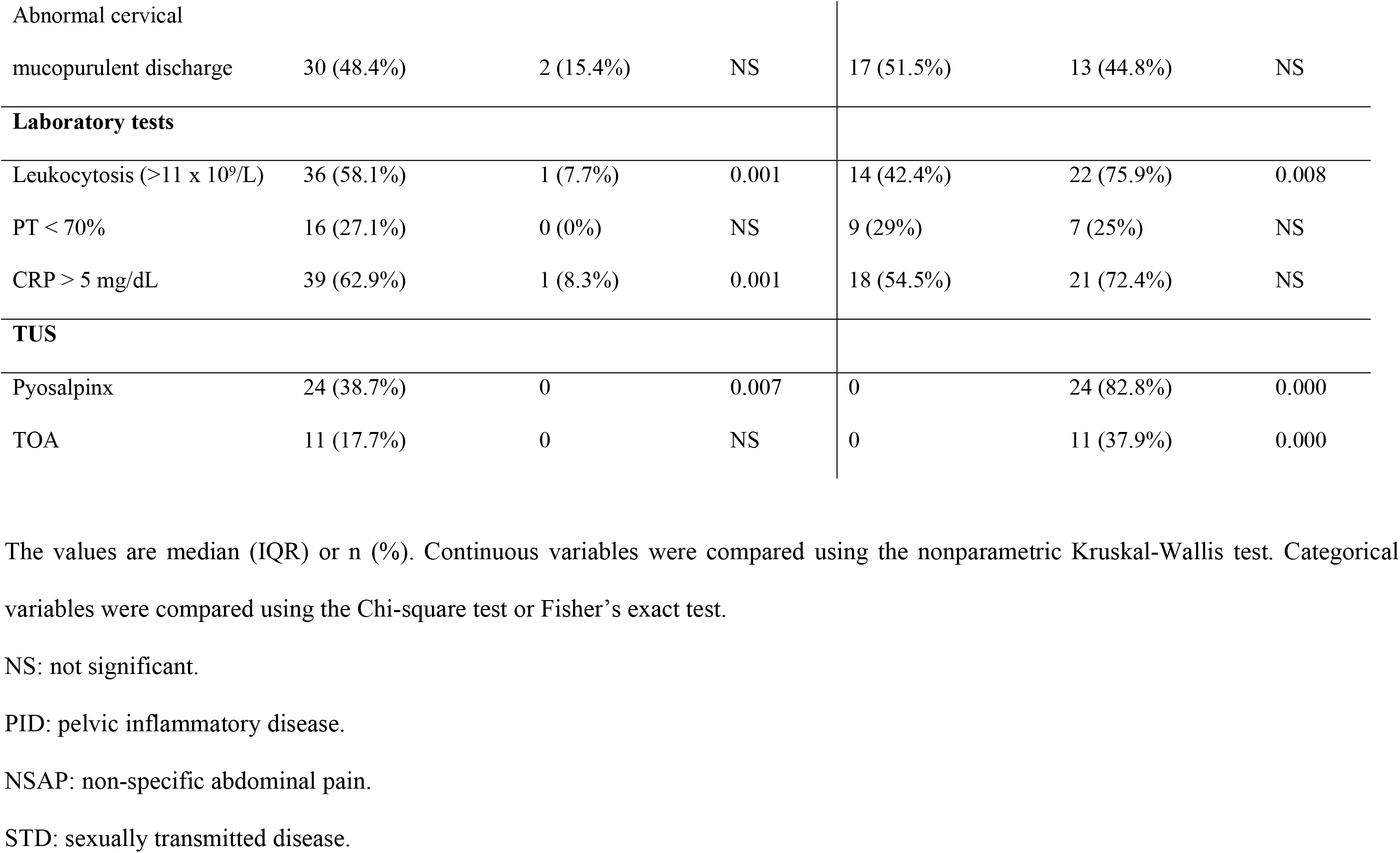
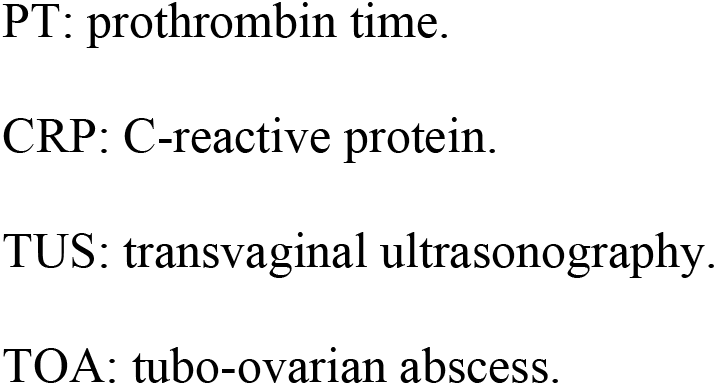
Comparion of baseline characteristics, findings at physical examination, laboratory tests, and transvaginal ultrasonography between patients with pelvic inflammatory disease (PID) *vs*. non-specific abdominal pain (NSAP), and between patients with mild-to-moderate PID *vs*. severe PID.

### Microbiological diagnosis

Table 2 shows the results of the microbiological tests performed in each group of patients. For statistical purposes, CT/NG test results were also classified as at least one positive (CT and/or NG) or all negative (for both CT and NG). Endocervical swabs and urine samples from all patients were tested for CT/NG. Regarding endocervical samples, 19 patients (19/75, 25.3%) presented infection by either CT (14/75, 18.7%) and/or NG (7/75, 9.3%) (two cases of co-infection). Urine failed to detect NG in one case with NG in the endocervical swab. Concordance between the endocervical and urine samples was 98.7% with a Cohen’s kappa coefficient of 0.93. Concordance between the Anyplex® CT/NG and GeneXpert® CT/NG assays was 100%. However, the mean time to results was significantly shorter for GeneXpert® CT/NG than for Anyplex®CT/NG: 2.22 hours *vs.* 24.37 hours, respectively (*p*<0.001).

**Table 2.**
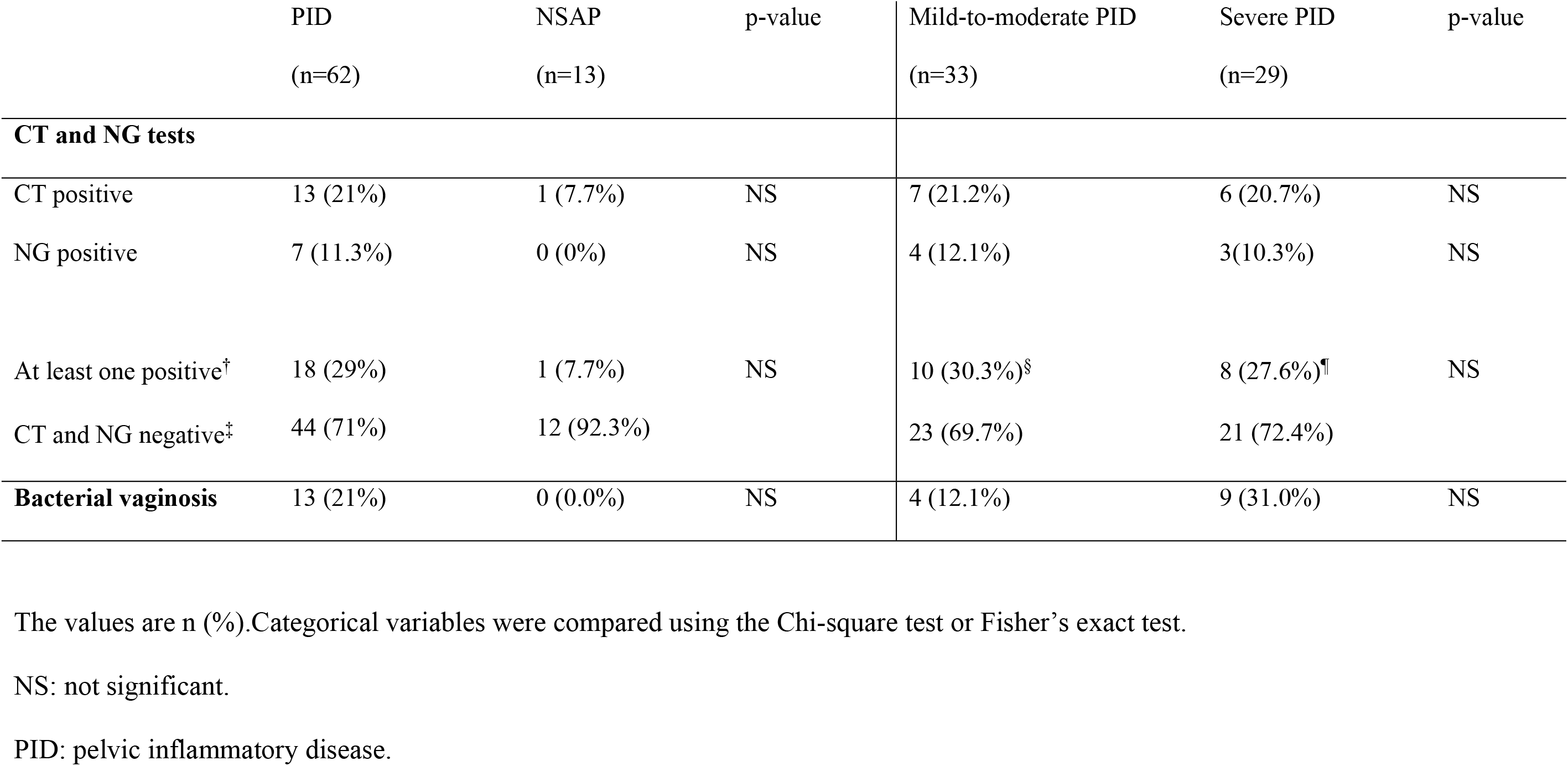
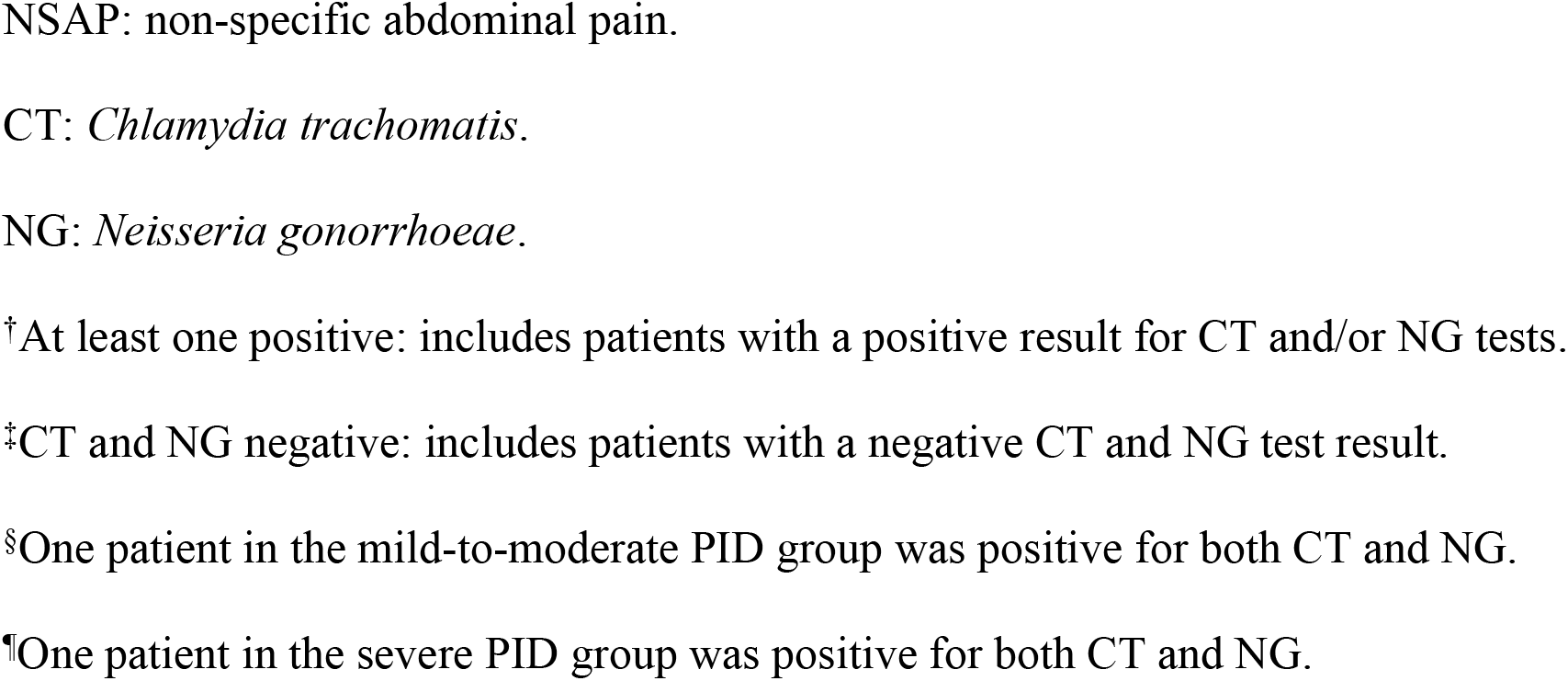
Comparison of microbiological results between patients with pelvic inflammatory disease (PID) *vs*. non-specific abdominal pain (NSAP) and between patients with mild-to-moderate PID *vs*. severe PID.

Urine cultures of all the patients were negative. Intraabdominal fluid was cultured in all patients requiring surgery (three with mild-to-moderate PID and six with severe PID), with four patients with severe PID being positive: 2 *E. coli*, 1 *Bacteroides fragilis* and 1 *Mycoplasma hominis*. Intraabdominal fluid obtained from patients who required surgery was also tested for CT/NG: two patients with mild-to-moderate PID were positive for CT and NG, respectively; and one patient with severe PID was positive for NG (who was also positive for *E. coli* at intraabdominal fluid culture).

No *Trichomonas vaginalis* was detected and only one case of *Mycoplasma genitalium* was detected in a patient with CT.

### Clinical management of patients and follow-up

Of 14 patients hospitalized with mild-to-moderate PID, three required surgery due to unsatisfactory evolution, with intraoperative findings of salpingitis confirming PID. Among the patients receiving outpatient treatment (n=19), one required hospitalization due to unsatisfactory evolution. The median hospital stay of the women with mild-to-moderate PID was 4.5 days (range: 1-16).

All patients with severe PID were hospitalized except one, who refused to be admitted. Six of these patients required surgical treatment. The median hospital stay in these patients was 6 days (range: 3 to 15).

Finally, among 13 patients with NSAP with no clinical suspicion of PID, one was positive for CT. Appropriate treatment was administered, and the 6-week follow-up test was negative.

All the patients were followed at 6 weeks after treatment initiation. At this time endocervical samples and urine were only obtained from patients with a previously positive CT and/or NG result. All samples were negative except for that of one asymptomatic woman, who was positive for CT. She had been classified in the mild-to-moderate PID group and had previously been positive for CT. Nonetheless, as this woman was a sex worker, reinfection was more likely than persistent infection.

### Cost analysis

Table 3 provides the total cost and the mean cost per patient classified by severity of the condition and the presence or absence of CT and/or NG.

**Table 3.**
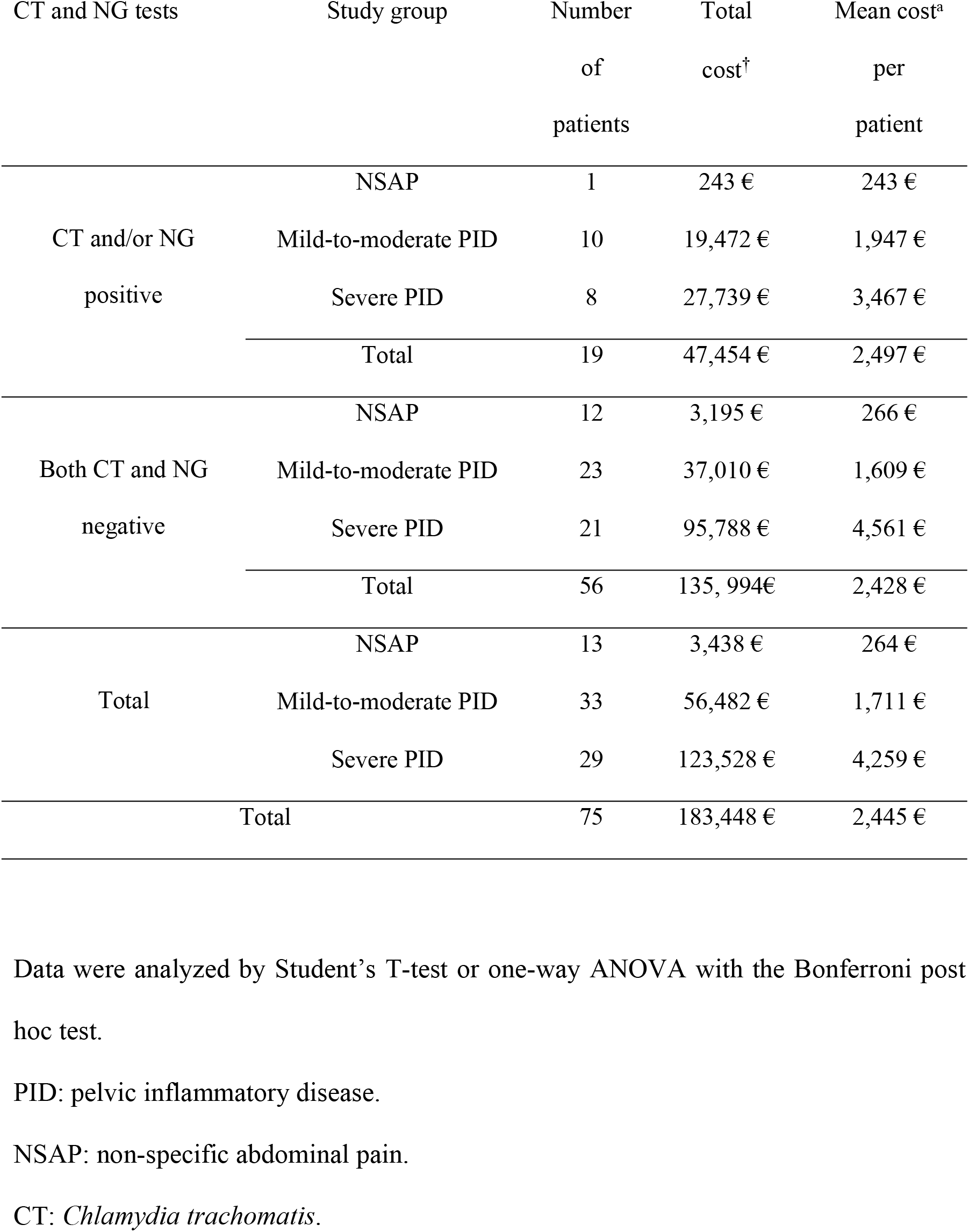
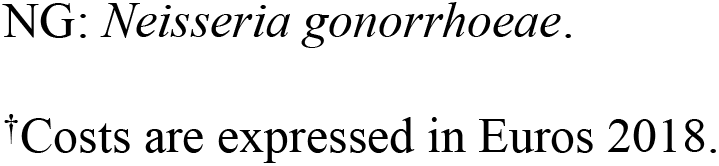
Total and mean costs per patient according to the study group and the result of the CT and NG tests.

There were significant differences in costs across severity levels but not between the presence or absence of CT and/or NG. Regarding infection severity, patients with severe PID presented the highest mean cost per patient, with the NSAP group showing the lowest mean cost per patient. It should be noted that surgery was more frequent among patients diagnosed with severe PID (20.7%) compared to those with mild-to-moderate PID (9%). This must have increased the mean cost per patient, especially among patients with severe PID who were negative for both CT and NG, five of whom required surgery (23.8%) compared with patients with severe PID with positive CT and/or NG results (12.5%).

## DISCUSSION

This study was designed to assess the presence of CT and/or NG infection in the lower genital tract of patients diagnosed with PID and NSAP, and to analyze the usefulness of a rapid molecular test to detect CT and NG for the diagnosis and clinical management of PID.

The diagnosis of PID is challenging^7,21^. According to the prevailing guidelines, all patients with suspected PID should undergo endocervical or vaginal tests for NG and CT, since a positive result supports its diagnosis^1,14,15^. However, in several studies no etiological agent was detected in approximately 65-75% of women with clinically diagnosed PID^6,7,9^. Thus, non-identification of the causal pathogen does not necessarily exclude the presence of PID^2,14,15^. In our study, no etiological agent was found in 71% of women diagnosed with PID, similar to previous reports^6,7,9,22^. Indeed, some authors have questioned whether this could represent a misdiagnosis of PID, especially in cases of mild-to-moderate PID^8,9^. Nonetheless, although the difficulty of case definition and diagnostic accuracy is a limitation for PID surveillance^7^. the present study followed the recommended diagnostic criteria for PID^2,14,15^.

Furthermore, we also included a group of patients with NSAP who did not meet the minimum criteria for suspicion of PID^14^ in order to establish if they could be misdiagnosed cases of subclinical PID. Of the 13 patients with NSAP, only one was positive for CT (7%). We were unable to establish whether this was a case of misdiagnosed mild PID or if the lower abdominal pain was attributable to other causes. Nevertheless, PID should be considered in all these cases^1^ taking into account that subclinical PID may be twice as common^1,21,23^ and that milder clinical manifestations of PID have increased as rates of NG have fallen^24^. Therefore, the presence of CT and NG in the lower genital tract should be evaluated in all sexually active women with STD risk factors who refer mild lower abdominal pain.

We also determined whether the use of a rapid molecular detection test could be helpful in the clinical management of patients diagnosed with PID. According to our data, concordance between rapid GeneXpert® CT/NG and standard Anyplex® CT/NG assays was 100%. However, results were more rapidly obtained with the rapid test. Despite of this fact, no differences were observed in the presence or absence of CT and/or NG in mild-to-moderate compared to severe PID. Moreover, the presence of CT and/or NG was not found to be a risk factor for a complicated clinical course (33% of patients undergoing surgery were positive for CT and/or NG compared to 30% of those not requiring surgery). Previous studies have reported similar data and recommend that PID management should be based on clinical features^9,25^. In the present study, hospitalization was decided according to clinical criteria^1,14,15,17^ and all patients were treated with broad-spectrum antibiotic regimens to cover likely pathogens, including CT and NG, irrespectively of the results^1,4,18^. Likewise, the economic analysis showed no cost differences between CT/NG-positive and negative patients. On the contrary, similar to previous reports^11,26^, patients diagnosed with severe PID presented the highest mean cost per patient due to the need for more complex treatments.

To our knowledge, this is the first study to assess the clinical utility of a rapid molecular test for CT and NG in patients diagnosed with PID. Nonetheless, our study had some limitations. The final sample size for the specific purpose of this study was 75 patients in a 12-month period obtained in a single hospital, where the study took place. It might seem a limitation but this sample is similar to previous studies^22^. The difficulty in obtaining a wider sample may be attributed to the fact that PID diagnosis is clinically difficult and its incidence is difficult to establish due to unspecific symptoms, subclinical cases and the possibility of misdiagnosis^1,7,8,21,23^. Other authors have also reported the difficulty in identifying and accurately diagnosing this condition^7^, which may be influenced by the health seeking behavior of patients, the clinical awareness of the attending ED physician and the results of complementary tests^7,8^. Therefore, although all the patients included were classified into the three groups according to recommended diagnostic criteria^2,14,15^,the possibility of a misdiagnosis cannot be ruled out, especially in cases of mild-to-moderate PID and NSAP. Another drawback is that the results from the rapid molecular test were not available for decision-taking since they were reported at the same time as those of the standard test. Nevertheless, as described previously, patient management should be based on clinical criteria, irrespectively of the presence or absence of CT and/or NG^1,2,7,14,15^.

Our study also has several strengths. Firstly, it was a prospective study and patients were classified into the three groups according to clinical criteria^2,14,15^. Another important issue is that two molecular methods were compared to detect CT and NG, with total concordance between both of them. However, rapid molecular tests (GeneXpert®) require little sample manipulation as extraction, amplification and detection occur in the cartridge allowing results to be obtained in a shorter time, and they may be used in settings with limited infrastructure to develop a standard molecular technique. Finally, we compared CT and NG detection in endocervical and urine samples in order to assess the sensitivity of both samples. NG was not detected in urine in a patient with NG in the endocervical swab, thus the sensitivity for urine was slightly lower than that of endocervical samples. This is concordant with previous reports^27^, and therefore, endocervical or vaginal samples but not urine should be used for CT and NG tests in women.

In conclusion, the use of a rapid molecular test for CT and NG in patients with clinical suspicion of PID and patients with NSAP could help in the differential diagnosis of abdominal pain in women at risk of STD in clinical settings in which a standard technique is not available. Moreover, the use of these tests, together with increased awareness among medical staff, might increase the diagnosis of mild or subclinical PID. Nonetheless, positive CT and/or NG test results may not be helpful for clinical management and the only cost difference relates to disease severity. Further research is warranted to assess these issues and to evaluate the usefulness of rapid molecular tests in clinical settings in which standard methods are not available.

## ACKNOWLEDGMENTS

This study was funded by the Hospital Clínic de Barcelona, Fundació Privada Clínic per a la Recerca Biomèdica (project number: CP041887), Barcelona Institute for Global Health (ISGlobal) and Cepheid Inc (Sunnyvale California, US). Cepheid Inc contributes economically to this project. Cepheid Inc did not participate in the study design, in the collection, analysis and interpretation of the data, in the writing of the report and in the decision to submit the paper for publication.

## DISCLOSURE

JM was contracted part time during six months in 2016 with funding obtained by Cepheid Inc.

